# Identification of Proteinase Activated Receptor (PAR) cleaving enzymes in human osteoarthritis knee joint synovial fluids

**DOI:** 10.1101/2020.10.21.336693

**Authors:** Arundhasa Chandrabalan, Andrew Firth, Robert B Litchfield, C Thomas Appleton, Alan Getgood, Rithwik Ramachandran

## Abstract

**Objective:** Osteoarthritis (OA) is the most prevalent joint disorder with incidence increasing worldwide. Mechanistic insights into OA pathophysiology are still evolving and there are currently no disease-modifying OA drugs available. It is well established that an increase in proteolytic enzyme activity is linked to progressive degradation of the cartilage in OA. Proteolytic enzymes can also trigger inflammation through activation of a family of G-protein coupled receptors (GPCRs) called the Proteinase Activated Receptors (PARs). Here we sought to characterize the PAR activating enzyme repertoire in human OA knee joint fluids.

**Methods:** Human knee joint synovial fluids derived from twenty-five OA patients and four healthy donors were screened for PAR cleavage activity using novel genetically encoded human PAR biosensor expressing cells. The class or type of enzymes cleaving the PARs was further characterized using enzyme-selective inhibitors and enzyme-specific fluorogenic substrates.

**Results:** Activity of PAR1, PAR2 and PAR4 activating enzymes were identified at substantially different levels in OA patients relative to healthy knee joint synovial fluids. Using enzyme class or type selective inhibitors and fluorogenic substrates we found that serine proteinases, including thrombin-like enzymes, trypsin-like enzymes, and matrix metalloproteinases are the major PAR activating enzymes present in the OA knee synovial fluids.

**Conclusions:** Multiple enzymes activating PAR1, PAR2 and PAR4 are present in OA joint fluids. PAR signalling can trigger pro-inflammatory responses and targeting PARs has been proposed as a therapeutic approach in OA. Knowledge of the PAR activators present in the human knee joint will guide study of relevant signaling events and enable future development of novel PAR targeted therapies for OA and other inflammatory joint diseases.

## INTRODUCTION

Osteoarthritis (OA) is the most common form of arthritis and a major health care burden worldwide with an estimated 250 million people currently affected worldwide^1–3^. In the context of this substantial global burden, most patients with OA receive symptomatic treatment alone given the paucity of effective disease-modifying agents^2^. This difficulty in identifying effective therapeutics stems in large part from the complex nature of this chronic disease, with many modifiable and non-modifiable risk factors including age, obesity, sex, trauma, physical activity, genetics and numerous environmental influences implicated in OA^3^. Irrespective of the etiology, a key feature of OA is the progressive degradation of the cartilage with an increase in proteolytic enzyme activity linked to this damage^4–6^. Synovial inflammation^7^ and bone remodelling^8^ are also important pathophysiological events in OA.

The role of proteolytic enzymes as important regulators of damage in OA has been long recognized. The matrix metalloproteinases (MMPs) in particular have received much attention with their collagenolytic activity playing an important role in OA pathology^9^. The enzymatic activity of MMPs and aggrecanases weaken the cartilage matrix, making it more susceptible to mechanical disruption during joint loading and movement. In addition to MMPs, enzymes from the coagulation, fibrinolytic and immune cells are also present in synovial fluids. Other enzymes such as the cysteine proteinase cathepsin-K can also degrade proteins in the cartilage and bone extracellular matrix (ECM)^4^. In addition to the key matrix-degrading MMPs and cysteine proteinases, serine proteinases such as matriptase, coagulation cascade enzymes, kallikrein-related peptidases and neutrophil enzymes, derived from both structural and immune cells in the joints, contribute to ECM remodelling, tissue healing, pain, inflammation and immunity, and as important regulators of OA pathogenesis^10^.

A key mechanism by which these proteolytic enzymes perpetuate pathological conditions in the joint is by activating a four-member family of G-protein coupled receptors (GPCRs) called the Proteinase Activated Receptors (PARs, PAR1-4)^11^. Expression of PAR1, PAR2, PAR3 and PAR4 are documented in cells of the synovium, cartilage, bone and neurons^12^. The role of PAR3 in physiology and pathophysiology however remains poorly understood since this receptor cannot signal independently^12,13^. PAR activation is generally considered to be proinflammatory^14–17^ and proalgesic^18^, though roles of specific receptors and associated signalling events remain unclear with model-specific and mechanistic differences in protection afforded by receptor deletion noted^19,20^.

A significant challenge in understanding the role of PARs in joint disease relates to the variety of enzymes that can activate these receptors^11,21^. PAR1 is classically described as thrombin activated receptor, but certain MMPs and neutrophil-derived enzymes including elastase, proteinase-3 and cathepsin-G also cleave and activate PAR1. PAR2 is similarly activated by a number of trypsin-like enzymes such as trypsin, matriptase, mast cell tryptase, mast cell chymase, neutrophil-derived enzymes^22^ and cathepsins^23^. PAR4 can be activated by thrombin, trypsin and neutrophil cathepsin-G^24,25^. Irrespective of the proteinases involved, receptor activation requires enzymatic cleavage of extracellular N-terminus of the receptor to unmask a receptor motif called the tethered-ligand, which then binds intramolecularly and activates the PARs. Interestingly, different enzymes cleave the PARs at different positions on the receptor N-terminus, leading to different tethered-ligands being generated, and different signalling cascades being turned on^26^. This diversity of PAR regulating enzymes present in the joint spaces is poorly understood and this remains a significant challenge in understanding PAR signalling in OA. Compounding this problem further, there exist species-specific differences in the complement of both proteinases and the PAR receptors^21,27^. In this context, previous studies have not directly examined PAR activation by proteinases present in human osteoarthritic knee joints. In the current study, we have sought to address this problem by using novel genetically encoded PAR biosensors to broadly classify synovial fluid enzymes in patients with knee OA that can activate PAR1, PAR2 and PAR4. We find that serine proteinases and metalloproteinases are present in OA joint fluids and are able to substantially activate PAR1 and PAR2 receptors, with more modest levels of PAR4 activation evident.

## MATERIALS AND METHODS

### Materials

Thrombin (human plasma, high activity, 5000 U, Calbiochem-EMD Millipore) and trypsin (porcine pancreas, Type IX-S, 13000-20000 BAEE units/mg protein, Sigma-Aldrich) stock solutions were made in 25 mM 4-(2-hydroxyethyl)-1-piperazine ethanesulfonic acid (HEPES, Fisher Scientific). The thrombin-selective inhibitor PPACK.2HCl and MMP Inhibitor V (ONO-4817) were obtained from Calbiochem (Millipore Sigma). The broad-spectrum MMP inhibitor batimastat (BB-94, ≥98%) was from Sigma-Aldrich. Soybean trypsin inhibitor (STI) was from Thermo Fisher Scientific or Millipore Sigma. The fluorogenic substrates, Bz-Phe-Val-Arg-AMC.HCl (Thrombin substrate III) and Boc-Gln-Ala-Arg-AMC.HCl (Trypsin substrate) were from Bachem, Suc-Ala-Ala-Pro-Phe-AMC (Chymotrypsin substrate II) and Z-Gly-Gly-Arg-AMC.HCl (Urokinase substrate III) were from Calbiochem, and MCA-Lys-Pro-Leu-Gly-Leu-Dpa(DNP)-Ala-Arg-NH2 (MMP substrate FS-6) was from Sigma-Aldrich. The stock solutions of the enzyme inhibitors and fluorogenic substrates were prepared according to manufacturer’s instructions. All samples were diluted to appropriate working concentrations in Hanks’ Balanced Salt Solution containing CaCl2 and MgCl2 (HBSS, Gibco ThermoFisher Scientific), except for the fluorogenic substrates which were diluted in Tris-NP40-calcium buffer [50 mM Tris.HCl pH 8 (Fisher BioReagents), 0.2% Nonidet P-40 Substitute (Roche) and 1.5 mM CaCl2 (Fisher BioReagents)].

### Patients criteria

Patients with a primary diagnosis of symptomatic knee OA undergoing a coronal plane alignment correction by opening wedge proximal tibial osteotomy were included. Patients undergoing concomitant procedures such as ligament reconstruction, meniscal transplantation or cartilage restoration were excluded. Patients with inflammatory arthropathy or past history of joint infection were also excluded. All patients completed a radiological examination including weight bearing anteroposterior, 45-degree flexed posteroanterior, lateral and standing hip-knee-ankle alignment views. The amount of radiographic knee OA was graded independently by two orthopedic surgical fellows on a scale of grade 0 (none), 1 (doubtful), 2 (mild), 3 (moderate) and 4 (severe) according to the Kellgren and Lawrence (KL) classification system^28^. Absence of knee symptoms and radiographically confirmed KL 0 was considered healthy. Patients also completed Knee Injury and Osteoarthritis Outcome Score (KOOS)^29^, the Western Ontario and McMaster Universities Arthritis Index (WOMAC)^30^ and Visual Analog Scale (VAS) pain score^31^ questionnaires at baseline prior to surgery. The total KOOS is a mean percentage score of the five subscales encompassing pain, symptoms, activities in daily living function (ADL), sport and recreation function (Sport/Rec) and quality of life (QOL), score mean^29,32^. The total WOMAC is a mean percentage score of the three subscales encompassing pain, stiffness and physical function^30^. In addition, each subscale score was calculated independently and transformed to a percentage score. Each score was converted to a percentage score by using the formula below, where KOOS and WOMAC transformed score of 100% represents no problems and 0% indicates extreme problems. VAS pain score rates patient’s pain at rest and during activity (move) on average over the past week, and the scale ranges from 0 (no pain) to 10 (worst pain possible). The mark placed along the scale (0-10 cm) by the patient is measured as the level of pain for each situation^31^.

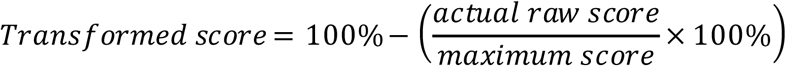

### Synovial Fluids

Healthy donor knee synovial fluids were obtained from Rheumatology Centre at St. Joseph’s Health Care London (C.T.A), under approved ethics ID # REB 109255. Synovial fluid samples from OA patients undergoing realignment osteotomy were obtained from the Fowler Kennedy Sport Medicine Clinic and University Hospital at London Health Sciences Centre, London, Ontario, under approved ethics ID # REB 108039 approved by the Western University Health Sciences Research Ethics Board. At surgery, OA patient synovial fluids from knee joints were collected by an orthopedic surgeon (A.G or R.B.L) prior to commencing arthroscopy and proximal tibial osteotomy. All samples were immediately placed on ice and transferred to the laboratory, centrifuged at 10000 × *g* for 1 minute, aliquoted and frozen at −80 °C. All joint fluids were tested for presence of any blood with the Fecal Occult Blood Test kit (Immunostics Hema-Screen) and the samples showing presence of blood were excluded from further analysis.

### Cell culture

Chinese Hamster Ovary (CHO-K1, Sigma) cells were cultured in Ham’s F-12 (1×) Nutrient Mix supplemented with 1 mM L-Glutamine, 100 U ml^-1^ penicillin, 100 µg ml^-1^ streptomycin, 1 mM sodium pyruvate, and 10% v/v heat-inactivated Fetal Bovine Serum (Gibco^®^ ThermoFisher Scientific). CHO cells stably transfected with the PAR biosensors cloned into pcDNA3.1(+) were cultured in complete F-12 medium with 600 µg ml^-1^ geneticin selective antibiotic (G418 Sulfate, Gibco^®^ ThermoFisher Scientific). The cells were grown in a T75 cell culture flask (Nunc) in a humidified cell culture incubator with 5% CO_2_ at 37 °C. Cells at ∼80-90% confluency were detached with phosphate-buffered saline (PBS, Gibco^®^ ThermoFisher Scientific) solution supplemented with 1 mM EDTA (Fisher Scientific), centrifuged at 180 × *g* for 5 minutes, and sub-cultured as appropriate.

### Cloning and Stable transfection

Human PAR2 with N-terminal nano-luciferase (nLuc, Promega) and C-terminal enhanced Yellow Fluorescent Protein (eYFP) tagged constructs have been previously described^33^. N-Terminal nLuc tagged human PAR1 and PAR4 constructs were similarly constructed by generating restriction enzyme sites for BspE1 and BamH1 by site-directed mutagenesis (QuickChange, Agilent technologies) and inserting nLuc in previously described eYFP tagged receptor constructs^34,35^. nLuc was located between the residues Glu^30^Ser^31^ in PAR1, Gly^28^Thr^29^ in PAR2, and Pro^23^Ser^24^ in PAR4. Fidelity of all constructs was verified by direct sequencing (London Regional Genomics Centre, Robarts Research Institute). CHO-K1 cells were stably transfected with the nLuc and eYFP tagged hPAR1, hPAR2 or hPAR4-pcDNA3.1(+) constructs by electroporation (Super Electroporator NEPA21 Type II, Nepa Gene) with 3 µg plasmid DNA and 1 × 10^6^ cells in 100 µl Opti-MEM (1×) Reduced Serum Medium (Gibco^®^ ThermoFisher Scientific). The electroporated cells were cultured in a non-selective complete medium in a 100 mm × 20 mm cell culture dish (Falcon, Corning) for 48-72 hours. The cells were subsequently maintained in G418 selective medium in a T75 flask for 7-14 days. G418-resistant cells expressing the construct (nLuc-hPAR1/2/4-eYFP) were clonally sorted by flow cytometry (FACSAriaIII, London Regional Flow Cytometry Facility, Robarts Research Institute) and expanded in G418 selective medium. The stably transfected reporter cell lines were characterized with a known hPAR agonist, thrombin on nLuc-hPAR1-eYFP, trypsin on nLuc-hPAR2-eYFP, and both thrombin and trypsin on nLuc-hPAR4-eYFP, using the luciferase assay technique described below. As a negative control, luminescence level in the parental CHO-K1 cells treated with PAR agonists thrombin and trypsin was also assessed.

### Luciferase assay

The presence of active PAR cleaving enzymes was measured by monitoring release of the N-terminal nLuc fusion tag. The nLuc-hPAR1/2/4-eYFP-CHO reporter cells were plated either in a 24-well or 96-well cell culture plates (polystyrene, flat-clear bottom, Nunclon Delta, Nunc, Thermo Fisher Scientific) at a cell density of 5 × 10^4^ cells per well in complete F-12 medium and cultured for 48 h. The cells were rinsed with HBSS (3 × 100 µl) and incubated with 100 µl HBSS at 37 °C for 15 minutes. 50 µl of cell supernatant from each well was transferred into a 96-microwell white plate (polystyrene, Nunclon Delta, Nunc, Thermo Fisher Scientific) and served as a measure of basal luminescence levels in each well. The biosensor expressing cells were then incubated with 50 µl of test samples, recombinant enzymes or controls, at 37 °C for 15 minutes. 50 µl of cell supernatant from each well was removed and transferred to a white plate as before. The Nano-Glo Luciferase Assay Substrate furimazine (2 µl ml^-1^, Promega) was added and the luminescence was measured on a luminometer (Mithras LB 940 Berthold Technologies plate reader, measurement time: 1 s per well).

### Fluorogenic substrate assay

Proteinase activity in synovial fluids was also measured using peptide-based fluorogenic substrates. The synovial fluid samples (10%) were either untreated or pretreated with enzyme inhibitors, PPACK (1 µM), STI (1 mg ml^-1^), BB-94 (10 µM) or ONO-4817 (10 µM) for 30 minutes at room temperature. Substrate cleavage was measured in a black clear bottom 96-well microplate (Greiner Bio-One) with the thrombin, trypsin, chymotrypsin, urokinase substrates (100 µM final concentration in 60 µl) and MMP substrate (25 µM final concentration in 60 µl). Immediately following the addition of the substrates, the fluorescence was measured every minute for 30 minutes on a fluorometer to obtain a kinetic curve. The substrates containing fluorescent 7-amino-4-methylcoumarin (AMC) were read on a PerkinElmer Victor plate reader (λ_Ex:Em_ 355:460 nm; measurement time: 0.1 s per well; lamp energy: 40000), and the MMP substrate containing the fluorophore (7-methoxy-coumarin-4-yl)acetyl (MCA) was read on a Cytation5 BioTek plate reader (fluorescence endpoint, λ_Ex:Em_ 325/20:390/20 nm; measurement time: 0.1 s per well; gain: extended; lamp energy: high, extended dynamic range).

### Graphical and statistical analyses

The luminescence values in the luciferase assay were normalized by subtracting the basal luminescence in HBSS treated samples. The concentration-effect curves for standard enzymes were plotted and analyzed using the standard slope (1.0) dose-response stimulation three parameters models with a non-linear regression curve fit in GraphPad Prism 8, and the values of logEC_50_ were obtained with their standard error of the mean (SEM). Each data point on a concentration-effect curve corresponds to the mean of at least three independent experiments (*N* ≥ 3), performed either in duplicates (*n* = 6) or triplicates (*n* = 9), with their SEM. The normalized luminescence measurements of the test samples were calculated as the percentage of the maximum response of the standard agonist, thrombin (3 U ml^-1^) for PAR1 and PAR4, and trypsin (100 nM) for PAR2. The data represents the mean of at least three independent experiments (*N* ≥ 3), performed in duplicates (*n* = 2), with their SEM. Mann-Whitney U test was utilized to compare any statistical significance between different groups, and Welch’s t-test was used to identify any statistical significance between untreated and treated groups at *p* < 0.05 or 0.01. Two-way analysis of variance (ANOVA) with Dunnett’s multiple comparisons test was utilized to determine any statistical significance at *p* < 0.05 relative to an untreated sample. The fluorescence measurements in the fluorogenic substrate assay were normalized by subtracting the fluorescence values obtained with the substrate alone (blank). The measurement at 10 minutes from the addition of substrate was used to express the enzyme activity obtained with synovial fluids. The data represents the mean of three independent experiments (*N* = 3), performed in duplicates (*n* = 2), with their SEM. Cohen’s kappa (*κ*) was used to determine the inter-rater agreement between the two clinicians when grading KL knee OA. Agreement was interpreted using the scale *κ* <0.20: slight agreement, *κ* = 0.21-0.40: fair agreement, *κ* = 0.41-0.60: moderate agreement, *κ* = 0.61-0.80: substantial agreement, and *κ* = 0.81-1.0: almost perfect agreement^36^. Pearson correlation coefficient analysis was used to determine any correlation between PAR activity and subscales of KOOS/WOMAC/VAS scores. Correlation analysis was performed using GraphPad Prism 8. Correlation coefficient was interpreted based on the reported scale,^37^ *r* = ±1.00 to ±0.90: a very strong correlation, *r* = ±0.89 to ±0.70: a strong correlation, *r* = ±0.69 to ±0.40: a moderate correlation, *r* = ±0.39 to ±0.10: a weak correlation, and *r* = ±0.10 to ±0.00: negligible correlation.

## RESULTS

### Patient characteristics

Demographics and radiographic data of the four healthy donors (**H1** - **H4**) and twenty-five OA patients (**1** - **25**) are presented in **Table 1**. Fifteen male and ten female patients ranging from doubtful/mild OA (KL grade 1) to moderate OA (KL grade 3) at the time of sample collection and healthy donors, one male and three females, with no knee symptoms or radiographic features of OA (KL grade 0) were included in the study. The two clinicians demonstrated moderate inter-rater agreement when grading KL knee OA (*κ* = 0.49, 95% confidence interval: 0.23 to 0.79). The mean age ± SD for the OA patients studied here was 46 ± 10 years and for healthy donors was 32 ± 15 years. Twenty of the twenty-five OA patients in this study fit the criteria for being overweight (BMI 25.0-29.9 kg m^-2^) or obese (BMI >29.9 kg m^-2^), whereas three of the four healthy donors were in the normal weight (BMI 18.5-24.9 kg m^-2^) category and one was overweight.

**Table 1.** Patients’ demographics and radiographic data.

| Patient | Sex | Age /<br>(years) | BMI (kg<br>m<sup>-2</sup>) | KL<br>grade | Patient | Sex | Age /<br>(years) | BMI (kg<br>m<sup>-2</sup>) | KL<br>grade |
| --- | --- | --- | --- | --- | --- | --- | --- | --- | --- |
| **H1** | M | 23 | 25.2 | 0 | **12** | M | 52 | 38.9 | 2 |
| **H2** | F | 22 | 24.6 | 0 | **13** | F | 42 | 37.8 | 2 |
| **H3** | F | 27 | 22.2 | 0 | **14** | F | 43 | 30.4 | 2 |
| **H4** | F | 54 | 22.1 | 0 | **15** | F | 44 | 24.4 | 2 |
| **1** | M | 24 | 24.0 | 1 | **16** | F | 53 | 35.1 | 2 |
| **2** | M | 38 | 29.5 | 1 | **17** | F | 55 | 34.8 | 2 |
| **3** | M | 46 | 34.3 | 1 | **18** | M | 49 | 24.8 | 3 |
| **4** | M | 49 | 32.2 | 1 | **19** | M | 50 | 29.4 | 3 |
| **5** | M | 53 | 34.3 | 1 | **20** | M | 52 | 34.1 | 3 |
| **6** | F | 27 | 30.9 | 1 | **21** | M | 54 | 25.9 | n.a. |
| **7** | F | 28 | 49.3 | 1 | **22** | M | 58 | 24.4 | 3 |
| **8** | F | 43 | 26.7 | 1 | **23** | M | 61 | 23.8 | 3 |
| **9** | M | 35 | 28.9 | 2 | **24** | F | 50 | 31.8 | 3 |
| **10** | M | 37 | 40.6 | 2 | **25** | F | 56 | 33.3 | 3 |
| **11** | M | 48 | 29.4 | 2 |  |  |  |  |  |

H, healthy donor; M, male; F, female; BMI, body mass index; KL, Kellgren and Lawrence system of knee OA classification; n.a., not available

### Characterization of PAR biosensor expressing cell lines

PAR biosensor expressing stable reporter cell lines nLuc-PAR1-eYFP-CHO, nLuc-PAR2-eYFP-CHO and nLuc-PAR4-eYFP-CHO were characterized with known PAR activating enzymes, thrombin and/or trypsin, by measuring luminescence in culture supernatants (**Figure 1**). The cleavage of PAR1, PAR2 and PAR4 by the canonical activator enzymes was assessed by measuring the release of the PAR N-terminal nLuc tag. Luminescence values were assessed in the presence of the nano-luciferase substrate furimazine. The obtained EC50 for thrombin (0.06 U ml^−1^, **Figure 1A**) on PAR1, trypsin (6 nM, **Figure 1B**) on PAR2, and thrombin (0.8 U ml^−1^, **Figure 1C(i)**) and trypsin (12 nM, **Figure 1C(ii)**) on PAR4 were consistent with the enzyme concentration range eliciting PAR signalling in previously published work^38–40^. As expected, parental CHO-K1 cells treated with thrombin and trypsin did not show any luminescence signal (data not shown) confirming the absence of artifact.

**Figure 1.**
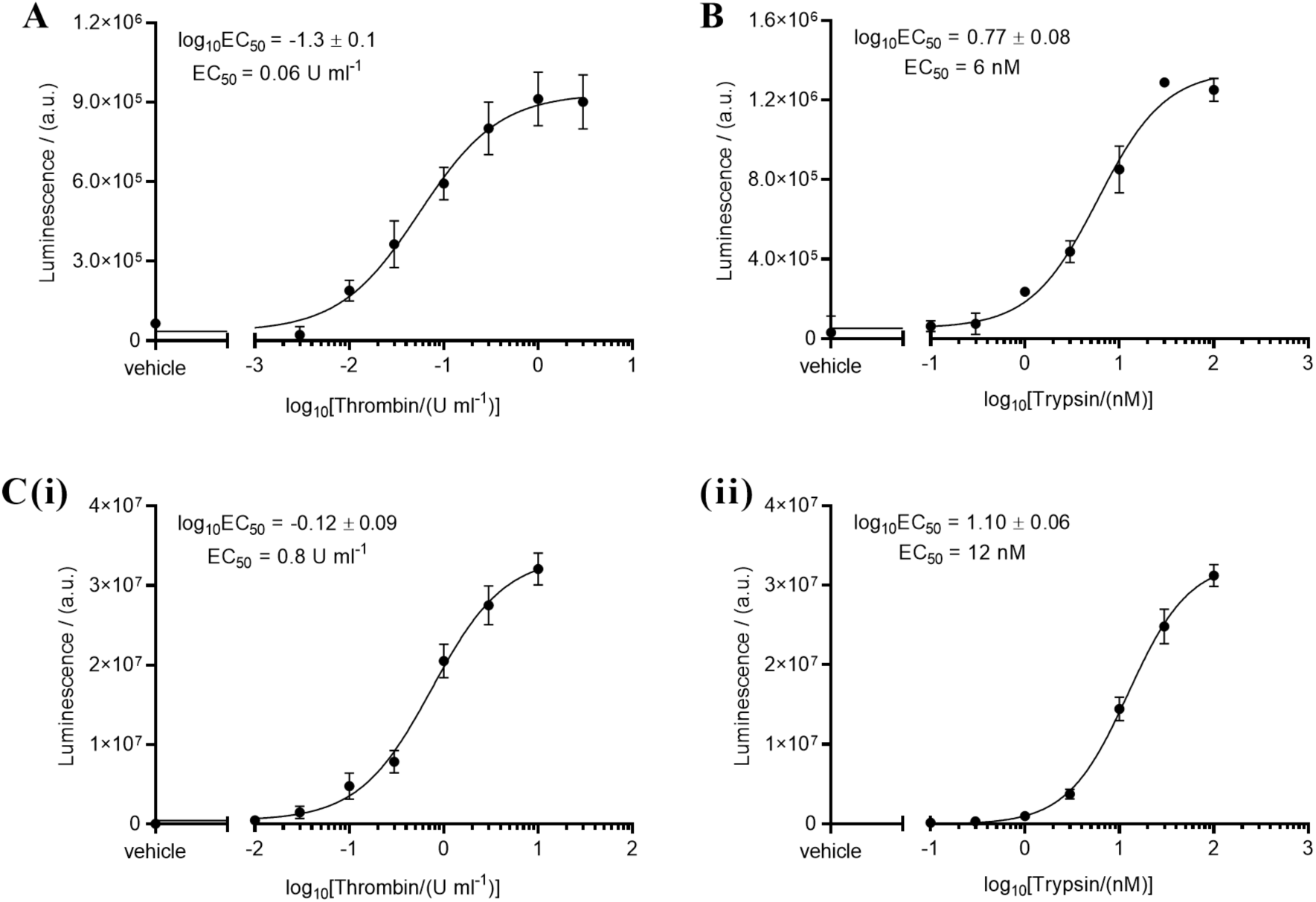
Characterization of PAR biosensor expressing reporter cell lines, (**A**) nLuc-PAR1-eYFP-CHO with thrombin (0.003 - 3 U ml^−1^), (**B**) nLuc-PAR2-eYFP-CHO with trypsin (0.1 - 100 nM), and (**C**) nLuc-PAR4-eYFP-CHO with (**i**) thrombin (0.001 - 10 U ml^−1^) and (**ii**) trypsin (0.1 - 100 nM), in half-log scale concentrations. Each data point on the concentration-effect curve represents the mean ± SEM (*N* ≥ 3).

### PAR1, PAR2 and PAR4 cleavage by enzymes in synovial fluids

Cleavage of PAR1, PAR2 and PAR4 was assessed with the four healthy (**H1 - H4**) and twenty-five OA patients (**1** - **25**) synovial fluids using the nLuc-hPAR1/2/4-eYFP-CHO reporter cell lines (**Figure 2**). Synovial fluid samples from five patients were contaminated with blood and were excluded from this study since these samples showed high enzyme activity which likely derived from the coagulation cascade. A high level of PAR1 cleavage was detected with all the patients (KL grades 1-3) synovial fluids relative to the healthy fluids (KL grade 0) examined (**Figure 2A, Supplementary Table S1**). When grouped by KL grade differences in PAR1 cleavage however failed to reach a statistically significant difference (*p* = 0.07 for healthy vs grade 2). Interestingly, cleavage of PAR2 (**Figure 2B, Supplementary Table S2**) and PAR4 (**Figure 2C, Supplementary Table S3**) were lower with the OA patients (KL grades 1-3) knee joint fluids compared to the healthy samples (KL grade 0). The difference between healthy and OA patients did not reach statistical significance for PAR2 cleavage, but the decrease in cleavage of PAR4 seen in OA patients was significantly lower (*p* < 0.05) than cleavage in healthy samples. There was no significant difference in activity observed between different OA grade (KL grade 1, 2 or 3) patients. It must be noted that these assays were done with a 1:10 dilution of the knee joint fluids and indicated cleavage of ∼10-20% of maximum thrombin (3 U ml^−1^) activity in PAR1 and ∼15-20% of maximum trypsin (100 nM) activity in PAR2, respectively. This would translate to the presence of ∼3-6 U ml^−1^ of thrombin-like enzymatic activity and 15-20 nM trypsin-like enzymatic activity in the knee joints.

**Figure 2.**
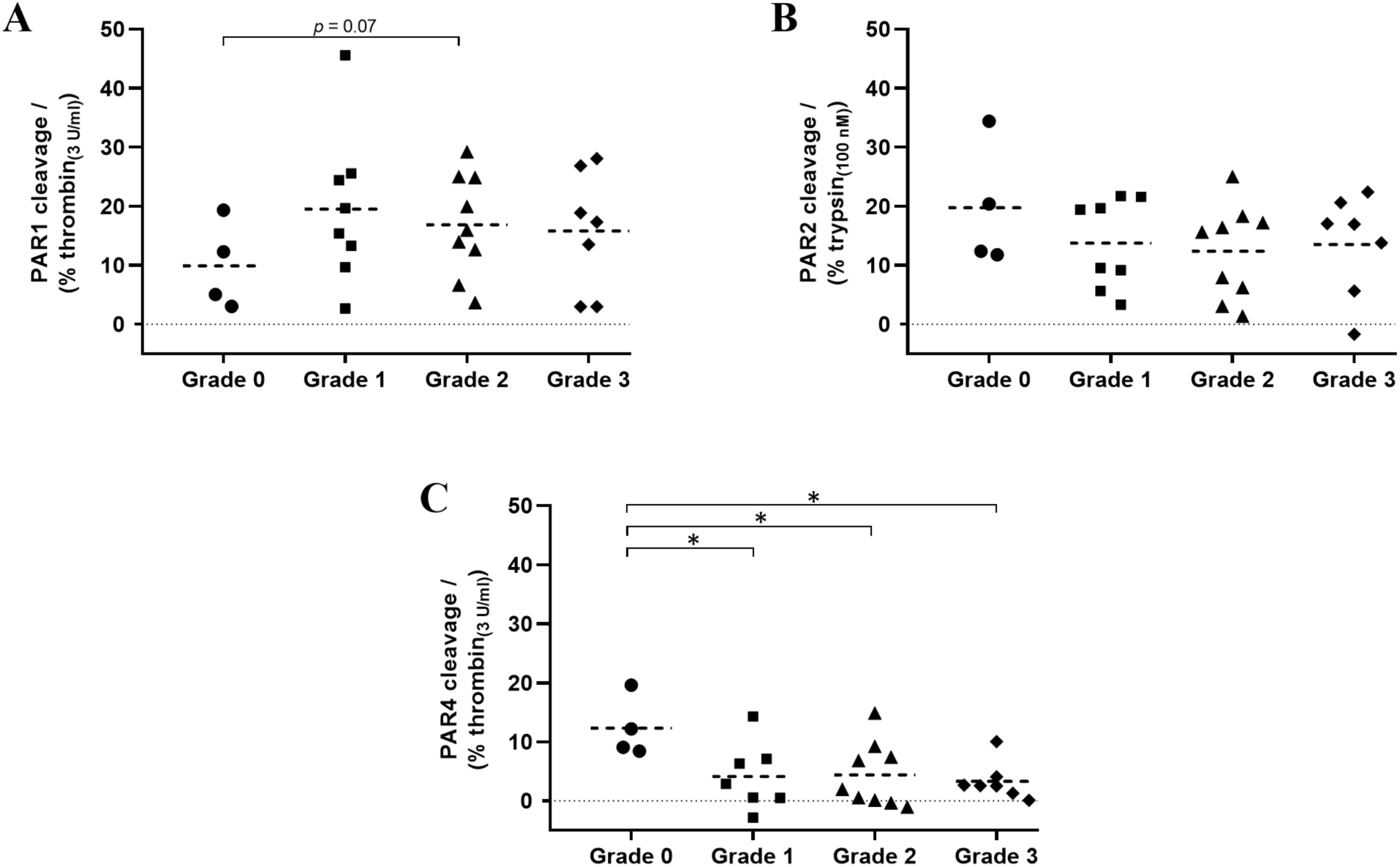
Cleavage of PARs by enzymes in human OA knee synovial fluids. Screening of healthy (Grade 0) and OA patients’ (Grade 1, 2 and 3) synovial fluids (10%) in nLuc-hPAR-eYFP-CHO cells for (**A**) PAR1, (**B**) PAR2, and (**C**) PAR4 cleavage. Thrombin (3 U ml^−1^) and trypsin (100 nM) were used as the PAR1/4 and PAR2 standard agonist, respectively. Mann-Whitney U test was utilized to compare any statistical significance between groups, \**p* < 0.05. Each data point on the scatter dot plot represents the mean ± SEM (*N* ≥ 3).

The observed PAR1, PAR2 and PAR4 cleavage were further compared against patients’ demographics including sex, BMI and age (**Figure S1**, Supplementary Information) to identify any associated trend in activity among the patients. Interestingly, PAR1 and PAR2 cleavage were significantly higher in males compared to females, while PAR4 cleavage was higher in females relative to males. There were no differences seen in PAR1/2/4 activity between normal weight (BMI 18.5-24.9 kg m^−2^), overweight (BMI 25.0-29.9 kg m^−2^) and obese (BMI >29.9 kg m^−2^) patients. PAR1 cleaving enzymes were insignificantly lower in synovial fluids from older adults (>55 years old) relative to young (18-35 years old) or middle-aged (36-55 years old) adults, whereas PAR4 cleavage was significantly low in middle-aged adults compared to young adults. Given the relatively small sample size in different categories, the trends observed here are interesting but need to be confirmed in a larger patient pool in future studies.

### Classification of PAR cleaving enzymes in synovial fluids

In order to understand the class/type of PAR cleaving enzymes present in the synovial fluids we used enzyme inhibitors broadly targeting serine proteinases that are known to activate PARs, such as thrombin-like (PPACK), trypsin-like (STI) enzymes, and MMPs (BB-94), and examined PAR biosensor cleavage by inhibitor pretreated synovial fluids in the reporter cell lines. PPACK is a potent, irreversible, thrombin selective inhibitor,^41,42^ which also inhibits coagulation factors VIIa and XIIa, tissue plasminogen activator (tPA), kallikrein,^43^ and urokinase^44^. STI is an inhibitor of trypsin^45^ that also inhibits chymotrypsin^45^, matriptase^46^, plasmin and plasma kallikrein to a lesser extent^47^. BB-94 is a potent, reversible, broad-spectrum MMP inhibitor targeting peptidases in families M10 and M12, including MMP-1, 2, 3, 7, 8, 9, 12, 13, 28, ADAM-8, 19, DEC1, and meprin-α, β^48–50^.

Due to limited volume of synovial fluids available, we could not examine any of the healthy fluids pretreated with inhibitors and only a subset of patient samples that were available in sufficient volumes (**1, 3, 5, 10, 17, 19, 20** and **23**) were assessed for PAR1 and PAR2 cleavage in the presence of enzyme inhibitors (**Figure 3**) and none of the samples were tested on PAR4 with inhibitors since initial experiments with patients fluids did not show significant cleavage of this receptor (**Figure 2C**). In the positive controls, cleavage of PAR1 by thrombin (3 U ml^−1^) was completely blocked by PPACK (1 µM, **Figure 3A**) and the PAR2 cleavage by trypsin (10 nM) was significantly blocked by STI (1 mg ml^−1^, **Figure 3B**). The patient synovial fluids (10%) pretreated separately with PPACK and BB-94 showed substantial reduction in PAR1 cleavage although a significant inhibition was seen only with some of the samples (**Figure 3A**). Similarly, with STI and BB-94 treated fluids, a significant drop in PAR2 cleavage was found with some of the samples (**Figure 3B**). Overall, these results indicate that different OA patient synovial fluids contain different serine proteinases and metalloproteinases that can potentially cleave the PARs expressed in the knee joint.

**Figure 3.**
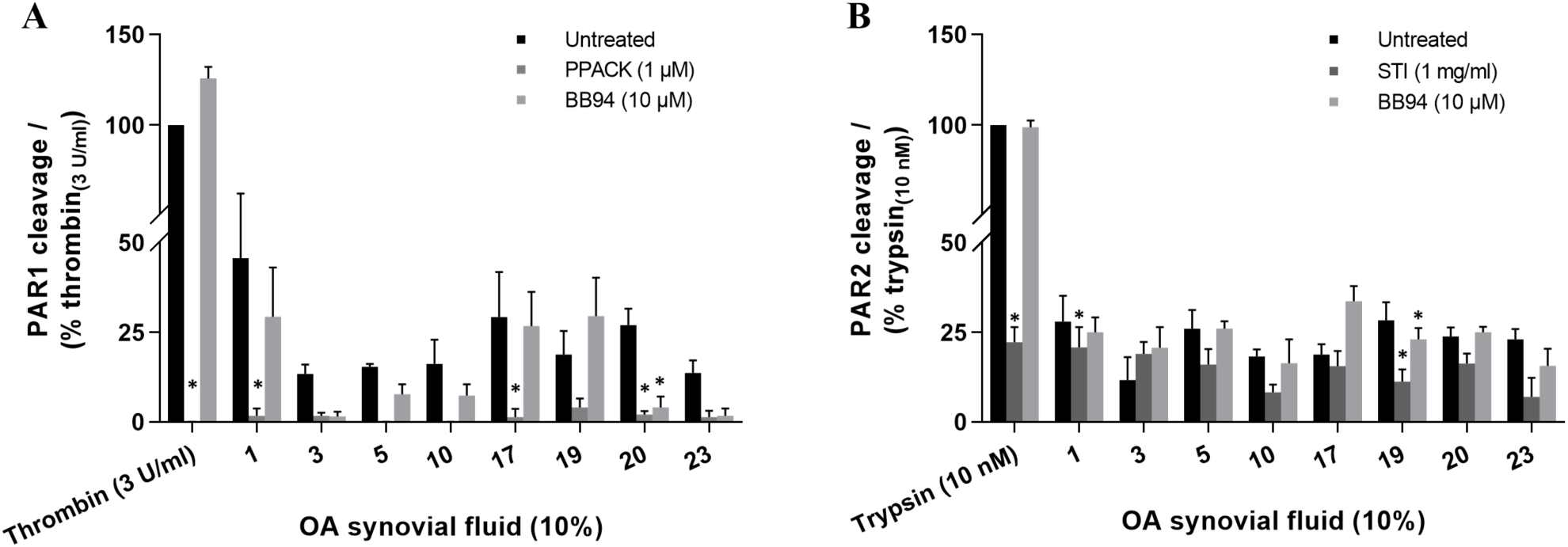
Classification of PAR cleaving enzymes in human OA knee synovial fluids. Screening of synovial fluids (10%) pretreated with an enzyme-selective inhibitor, PPACK, STI or BB-94, in nLuc-hPAR-eYFP-CHO cells for (**A**) PAR1 and (**B**) PAR2 cleavage. Two-way ANOVA with Dunnett’s multiple comparisons test was utilized to determine statistical significance at \**p* < 0.05 relative to the untreated sample response. Each bar represents the mean ± SEM (*N* ≥ 3).

As there appeared to be more than one class/type of enzymes present in the synovial fluid that cleave PARs, a complete inhibition of PAR1/2 cleavage with a single enzyme inhibitor is not likely to occur. We therefore used peptide-based fluorogenic substrates of thrombin, trypsin, chymotrypsin, urokinase, and MMPs to clarify the nature of enzymes present in the synovial fluids (**Figure 4**). With the use of thrombin substrate Bz-Phe-Val-Arg-AMC, trypsin and matriptase substrate Boc-Gln-Ala-Arg-AMC, and a broad-spectrum MMP substrate MCA-Lys-Pro-Leu-Gly-Leu-Dpa(DNP)-Ala-Arg-NH2 which can be hydrolyzed by MMP-1, 2, 3, 7, 8, 9, 12, 13, 14, 16, 20, ADAM-10, 17/TACE and BACE2^51– 54^, significant levels of thrombin-like enzymes (**Figure 4A**), trypsin-like enzymes (**Figure 4B**) and MMP (**Figure 4C**) activity were detected in the twenty-five patients 10% synovial fluids (KL grades 1-3) tested. However, no cleavage of the chymotrypsin sensitive substrate Suc-Ala-Ala-Pro-Phe-AMC, and the urokinase and plasminogen activators sensitive substrates Z-Gly-Gly-Arg-AMC.HCl was detected in any of the twenty-five patient synovial fluids (data not shown). Remarkably, thrombin-like and trypsin-like enzymes were significantly higher in different OA grade samples compared to healthy synovial fluids (**Figure 4A**,**B**), which was consistent with the high PAR1 cleavage found with the OA patient samples in the PAR biosensor assay (**Figure 2A**). Due to inadequate volume of healthy samples (KL grade 0), we were not able to test the healthy fluids using the MMP substrate or the enzyme inhibitors and compare against the OA patient fluids (KL grades 1-3) activity. As observed in the PAR biosensor cleavage assay (**Figure 3**), a significant inhibition of thrombin-like (**Figure 4A**), trypsin-like (**Figure 4B**) and MMPs (**Figure 4C**) enzymes activity were found in all of the OA patient synovial fluids (KL grades 1-3) pretreated with PPACK (1 µM), STI (1 mg ml^−1^) or BB-94 (10 µM), respectively. In addition, here we used another MMP inhibitor ONO-4817 that selectively inhibits only MMPs in family M10 (MMP-1, 2, 3, 7, 8, 9, 12 and 13)^48,55^ to narrow down the list of MMPs involved in OA. As seen with the broad-spectrum MMP inhibitor BB-94, a significant reduction in MMP activity was seen in the ONO-4817 (10 µM) pretreated patients synovial fluids (**Figure 4C**). Together this set of experiments further confirmed the presence of significant levels of serine proteinases and metalloproteinases in the OA knee joint fluids.

**Figure 4.**
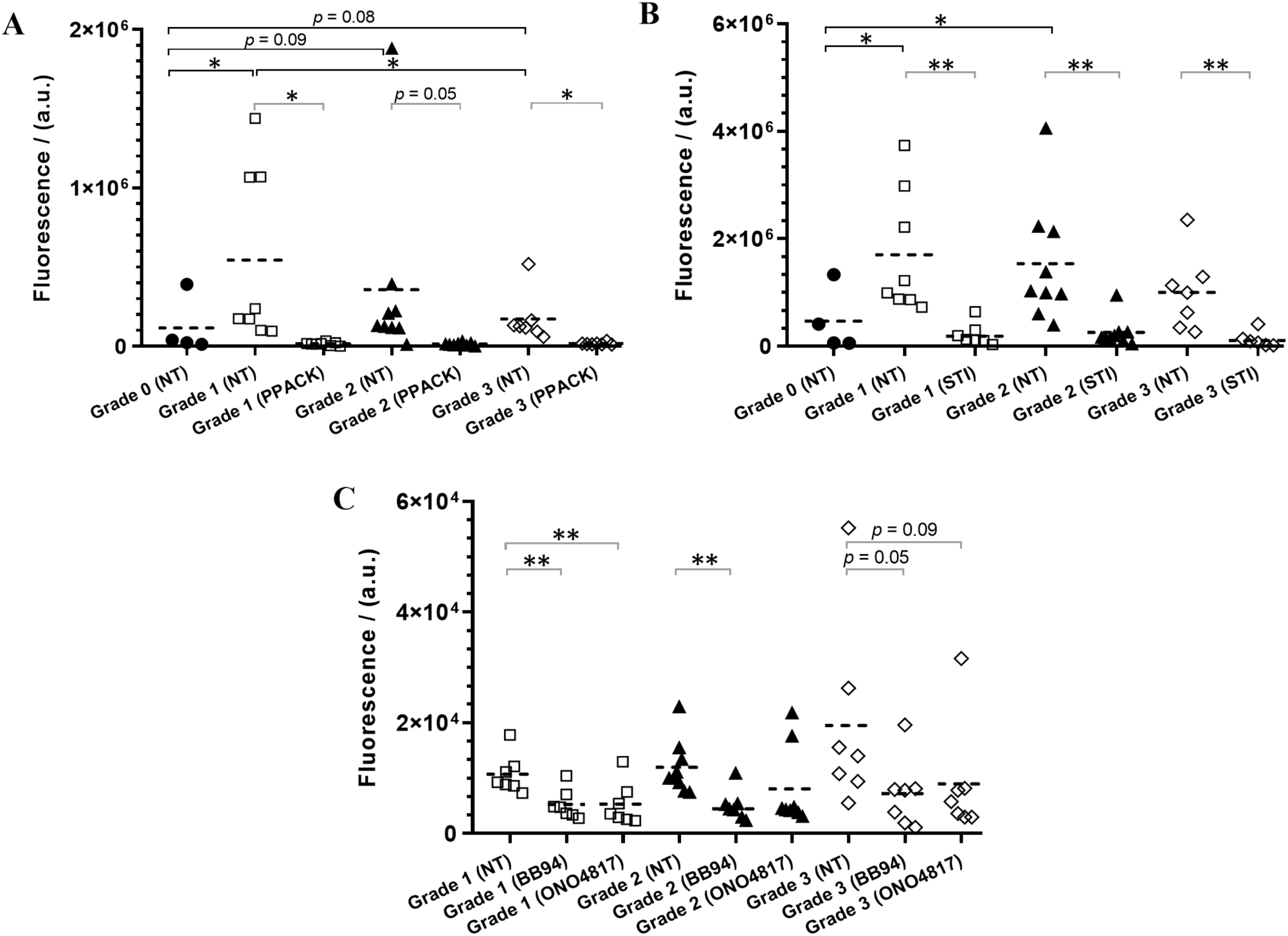
Identification of class/type of enzymes present in human OA knee synovial fluids. Cleavage of fluorogenic substrates by enzymes in healthy (Grade 0) and patients’ (Grade 1, 2 and 3) synovial fluids (10%), non-treated (NT) or pretreated with an enzyme inhibitor (PPACK, STI, BB-94 or ONO-4817), were measured using fluorogenic peptides, (**A**) Bz-FVR-AMC (thrombin-like enzymes), (**B**) Boc-QAR-AMC (trypsin-like enzymes), and (**C**) MCA-KPLGL-Dpa(DNP)-AR-NH2 (MMPs). Mann-Whitney U test was utilized to compare any statistical significance between different groups, and Welch’s t-test was used to identify any statistical significance between untreated and treated groups, \**p* < 0.05 and \*\**p* < 0.01. Each data point on the scatter dot plot represents the mean ± SEM (*N* ≥ 3), except the Grade 0 healthy samples data represents *N* = 1. Healthy samples were not examined with MMP substrate due to inadequate samples.

The substrate cleavage enzyme activity data was also analyzed in the context of the OA patients’ demographics (**Figure S2**, Supplementary Information). However, there was no substantial difference found in enzyme activity by sex of the patients in contrast to the trend observed in the PAR biosensor cleavage assay. As seen with PAR cleavage, enzyme activity did not show significant difference between normal weight, overweight or obese patients. It was however interesting to see a significantly lower level of thrombin-like enzyme activity in older adults relative to middle-aged adults as was observed with the thrombin receptor PAR1 cleavage (**Figure S1**). There was no significant difference seen in the trypsin-like enzyme activity between different age patients similar to that observed with the trypsin receptor PAR2 cleavage (**Figure S1**). More interestingly, MMPs activity were substantially higher in older adults compared to young or middle-aged adult patients. Together, there seems to be an interesting link existing between OA patient demographics, enzyme activity and PAR cleavage, though the sample size in this human pilot study was not large enough to make a strong statistical comparison.

In order to estimate the level of thrombin-like and trypsin-like enzymes in the patients’ synovial fluids, a kinetic standard curve of the fluorogenic substrate hydrolysis, Bz-Phe-Val-Arg-AMC and Boc-Gln-Ala-Arg-AMC by thrombin and trypsin, respectively was generated (**Figure 5**). From the time of substrate addition, a concentration-dependent enzyme-substrate activity was clearly observable at 10 minutes. Therefore, the fluorescence measurements at 10 minutes were used to compare the level of enzyme activity. Through interpolation of the measured enzyme activity in the thrombin and trypsin kinetic curves, it was found that concentrations of 0 - 0.01 U ml^−1^ thrombin-like and 0 - 0.3 nM trypsin-like enzymes are present in the 10% OA knee joint synovial fluids (KL grades 1-3) examined.

**Figure 5.**
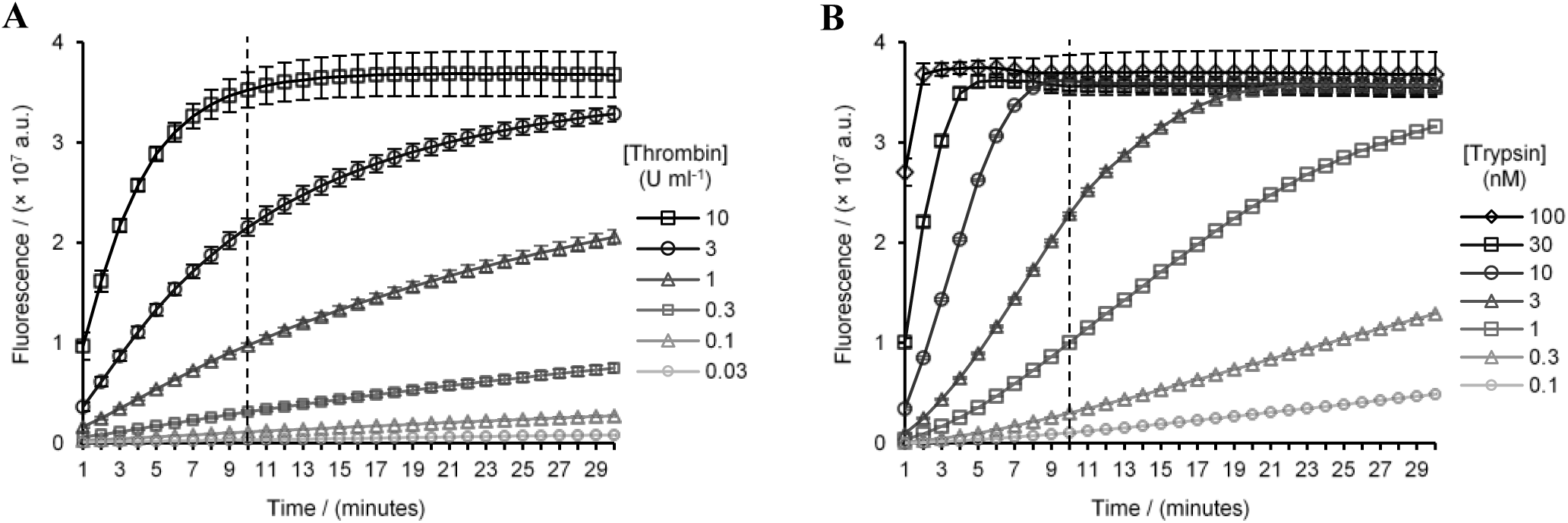
Kinetic curves of the thrombin and trypsin fluorogenic peptide substrate hydrolysis, (**A**) Bz-FVR-AMC (100 µM) by thrombin (0.03 - 10 U ml^−1^), and (**B**) Boc-QAR-AMC (100 µM) by trypsin (0.1 - 100 nM). Each data point on the curve represents the mean ± SEM (*N* = 3).

### Correlation of KOOS, WOMAC and VAS scores to PAR1/2/4 activity

OA patients self-reported arthritic questionnaires scores KOOS, WOMAC and VAS (Supplementary **Table S4**) were correlated to the obtained PAR1, PAR2 and PAR4 activities to identify if any correlation exists (**Figure S3**). Based on the Pearson correlation coefficient interpretation scale described under Methods and Materials section, a weak positive correlation was observed with all of the KOOS [*r*_Pain_ = 0.30, *r*_Symptom_ = 0.17, *r*_ADL_ = 0.17, *r*_Sport/Rec_ = 0.24, *r*_QOL_ = 0.34 (**Figure S3-A**)] and WOMAC [*r*_Pain_ = 0.28, *r*_Physical function_ = 0.17, *r*_Stiffness_ = 0.14 (**Figure S3-B**)] and VAS [*r*_Move_ = 0.25 (**Figure S3-C**)] scores against the PAR1 activity though these were not statistically significant. At rest the VAS score had a weak negative correlation *r*Rest = −0.15 against PAR1 activity. However, against the PAR2 activity, there was a weak positive correlation found only with one subcategory of the KOOS (*r*_QOL_ = 0.11), WOMAC (*r*_Pain_ = 0.13), and VAS (*r*_Move_ = 0.24) score, and a weak negative correlation was found with one of the KOOS (*r*_Symptom_ = −0.21) and WOMAC (*r*_Stiffness_ = −0.15) score, while all other scores had negligible correlation (**Figure S3**). As observed with the PAR1 correlation, most of the scores KOOS (*r*_Pain_ = 0.17, *r*_ADL_ = 0.20) and WOMAC (*r*_Pain_ = 0.22, *r*_Physical function_ = 0.20) had a weak positive correlation against the PAR4 activity although two of the KOOS (*r*_Sport/Rec_ = −0.10, *r*_QOL_ = −0.18) had a weak negative correlation and other subcategories and VAS scores had negligible correlation (**Figure S3**).

## DISCUSSION

Using novel biosensor expressing reporter cells and fluorogenic substrates we have monitored PAR receptor cleavage in knee joint fluids from twenty-five OA patients with disease severity ranging from KL grade 1-3. We find multiple active PAR1, PAR2 and PAR4 activating enzymes including serine proteinases and metalloproteinases in human knee joint synovial fluids tested. Curiously, levels of PAR1 activating enzymes were elevated and PAR4 activating enzyme levels decreased in the OA joint fluids but not in the healthy joint fluids studied here. This decline in PAR4 activating enzymes in OA joint fluids is interesting since some PAR1 and PAR2 activating enzymes, including thrombin and trypsin, also activate PAR4. This divergent activation of PAR1 and PAR4 in OA suggests that there are distinct PAR1 and PAR4 activating enzymes in the knee joint that should be identified in order to better understand roles of these receptors in OA pathogenesis.

Proteolytic enzymes are important mediators of joint health and pathophysiology. The role of MMPs in particular, as a class of enzymes that digest ECM components and cause cartilage degradation, is well established^56^. More recently it has emerged that other classes of enzymes such as serine proteinases are also upregulated in OA and may participate in proteolytic cascades leading to cartilage destruction^57^. All of the enzymes that are implicated in OA pathology also trigger pro-inflammatory signalling cascades through the activation of the PAR family of GPCRs^10^.

PARs have been studied in the context of arthritis using a number of animal models. PAR1 deficient mice showed decreased cartilage degradation and lower levels of synovial cytokine mRNA and MMP-13 mRNA in a model of antigen-induced arthritis^20^. PAR1 deficiency is also protective in a model of psoriatic arthritis driven by increased dermal expression of kallikrein 6^58^. In contrast, PAR1 deficiency did not afford significant protection in the destabilization of the medial meniscus (DMM) model of OA^19^. PAR2 deletion, or pharmacological inhibition, on the other hand has shown more consistent protective effects across multiple models, though there are some variations in the degree of reduction in cartilage erosion and subchondral bone thickening reported^15,19,59–61^. The role of PAR4 expressed in joint cells or tissues is yet unknown, however PAR4 activation significantly inhibits PAR2 agonist and transient receptor potential vanilloid-4 (TRPV4) agonist mediated visceral pain^62^. PAR4 activation can also decrease excitability in dorsal root ganglion neurons ^63,64^.

These disparate results across different models point to PAR signalling in different immune and joint cells contributing to the disease with PAR expression well established in chondrocytes, fibroblast-like synoviocytes, osteoblasts and immune cells^10,12^. Further studies with tissue-specific deletion of PARs across multiple models of arthritis is required to fully understand relative contributions of each receptor and cell type.

A number of known proteolytic enzyme activators of PARs are nonetheless implicated in OA and similar protection of joints in OA is also reported when the key PAR1 and PAR2 activating enzymes are inhibited. Serine proteinases involved in the coagulation and fibrinolysis cascades including thrombin, plasminogen activators and plasmin are well established as activators of PAR1 and show substantial increase in the inflamed joint of OA patients and in animal models^5,65,66^. MMPs, including MMP-1, 2, 3, 9, 13 and 14 are secreted in response to inflammatory cytokines and growth factors by chondrocytes and synoviocytes^6,56,67–69^. A number of these MMPs including MMP-1^70,71^, 2^72^, 3, 8, 9^73^,12^74^ and 13^75,76^ are able to activate PARs. In addition, immune cell derived proteinases such as mast cell tryptase^77^, neutrophil elastase^22,78^, proteinase-3^78^ and cathepsins^23,24^ are also able to cleave and activate PAR1 and PAR2. In this regard, our finding that multiple OA joint fluid enzymes can cleave PARs is particularly relevant. Firstly, this finding suggests that specific inhibition of individual enzymes may not be useful in treatment of OA since other enzymes could still perpetuate inflammatory signalling through these receptors. Pharmacological inhibition of individual receptors may instead be more beneficial. Secondly, in recent years it has emerged that not all enzymes trigger identical signalling responses through a PAR receptor, a concept called biased signalling^11,26,79^ that is now widely seen across multiple GPCRs. In addition to the canonical activation site of PARs [thrombin activation of PAR1 (cleavage at Arg^41^/Ser^42^) or PAR4 (cleavage at Arg^47^/Gly^48^) and trypsin activation of PAR2 (cleavage at Arg^36^/Ser^37^)], PAR cleavage by other enzymes occurs at various sites on the receptor N-terminus to reveal tethered-ligands and activate distinct signalling responses^13^. For example, cleavage of PAR1 by MMP (Asp^39^/Pro^40^, Leu^44^/Leu^45^, Phe^87^/Ile^88^), neutrophil elastase (Ala^36^/Thr^37^, Val^72^/Ser^73^, Arg^86^/Phe^87^) and proteinase-3 (Aal^36^/Thr^37^, Pro^48^/Asn^49^, Val^72^/Ser^73^, Ala^92^/Ser^93^) is reported to occur at multiple sites. In the case of PAR2, neutrophil elastase (Ala^66^/Ser^67^, Ser^67^/Val^68^)^22^ and cathepsin-S (Gly^40^/Lys^41^)^23^ also cleave the receptor at different sites than PAR1. Each of these different cleavage events can result in different signalling response, some of which may be protective. While we show that PARs can be cleaved by various enzymes in OA joint fluids, a thorough characterisation of receptor coupling to different G-protein and β-arrestin mediated signaling pathways is necessary to fully understand the role of this signaling system in joint health and pathology. The diversity of PAR activators in the joints also highlight the importance of understanding signalling differences elicited by the different PAR activating enzymes in testing appropriate therapeutic interventions. The significant decrease in PAR4 activating enzyme levels in OA patients compared to healthy controls could also be biologically significant and suggests the intriguing possibility that PAR4 is a mediator of analgesia in the knee joint that is lost in OA. It will also be important to test whether loss of PAR4 signalling is harmful in OA and if restoring PAR4 and inhibiting PAR1 can be protective.

This study was limited by a somewhat smaller sample size and our results could also be confounded by age (most healthy donors were younger). Our exploratory analysis of correlations with patients’ outcome could not therefore draw any strong conclusions. PARs however have crucial and well-established roles as regulators of inflammation and algesia and our studies advance our understanding of regulators in this important signalling system.

## CONCLUSIONS

We have established and characterized here a novel biosensor expressing cell line and an assay that allows rapid and facile screening of PAR activating enzymes from complex biological fluids. In a small population of OA patients, we were able to identify multiple enzyme types that target both PAR1 and PAR2. These studies broadly define proteinase classes that are present in the arthritic joint and guide future studies aimed at isolating and further characterizing these proteinases in the healthy and diseased joints.

## Authors Contributions

A.C., A.G. and R.R. designed the study overall. A.C. performed the experiments and wrote the initial draft of the manuscript with R.R. A.C., A.G. and R.R. contributed to the interpretation of the data. A.G. and R.B.L obtained synovial fluids from OA patients. T.C.A. obtained synovial fluids from healthy donors. A.F. performed statistical analysis on patients’ clinical data. The final manuscript was reviewed and approved by all authors.

## Acknowledgements

We thank Ms. Ashley Martindale and Ms. Stacey Wanlin for assistance with obtaining the patients’ synovial fluids, demographics/radiographic data, and arthritic questionnaire scores. We also thank Dr. Rajeshwar Sidhu and Dr. Taher Abdelrahman for helping grade patient radiographs. This study was supported by a catalyst grant from the Bone and Joint Institute at Western (R.R. and A.G.) and a Canadian Institutes of Health Research (CIHR) Project grant # 376560 (R.R).

---

**Figure S1.**
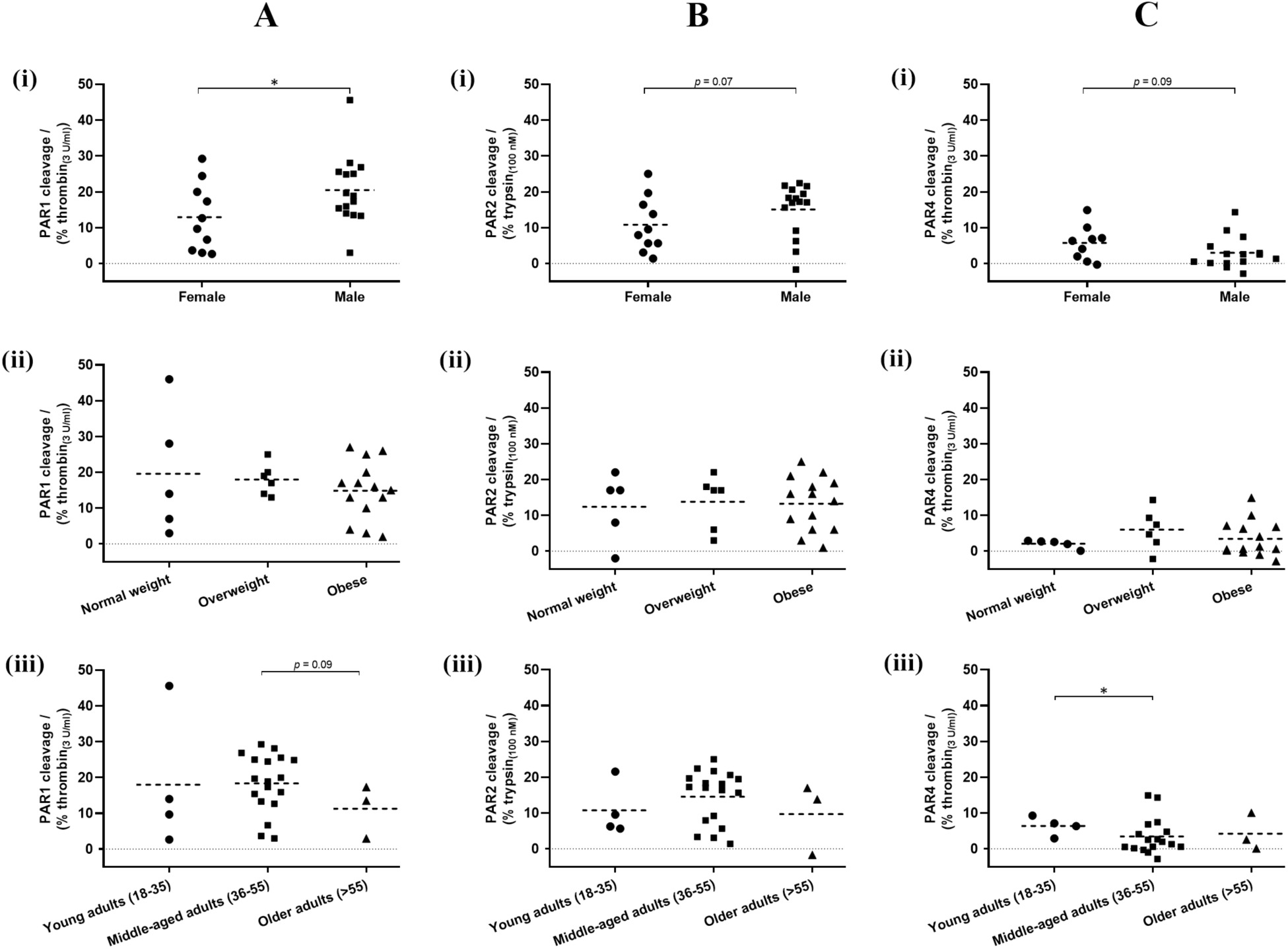
Cleavage of PAR1 (panel **A**), PAR2 (panel **B**) and PAR4 (panel **C**) observed with 10% patients’ synovial fluids in nLuc-hPAR-eYFP-CHO cells were plotted as a function of patient demographics, **(i)** sex, **(ii)** BMI and **(iii)** age. Mann-Whitney U test was utilized to compare any statistical significance between groups, \**p* < 0.05. Each data point on the scatter dot plot represents the mean ± SEM (*N* ≥ 3).

**Figure S2.**
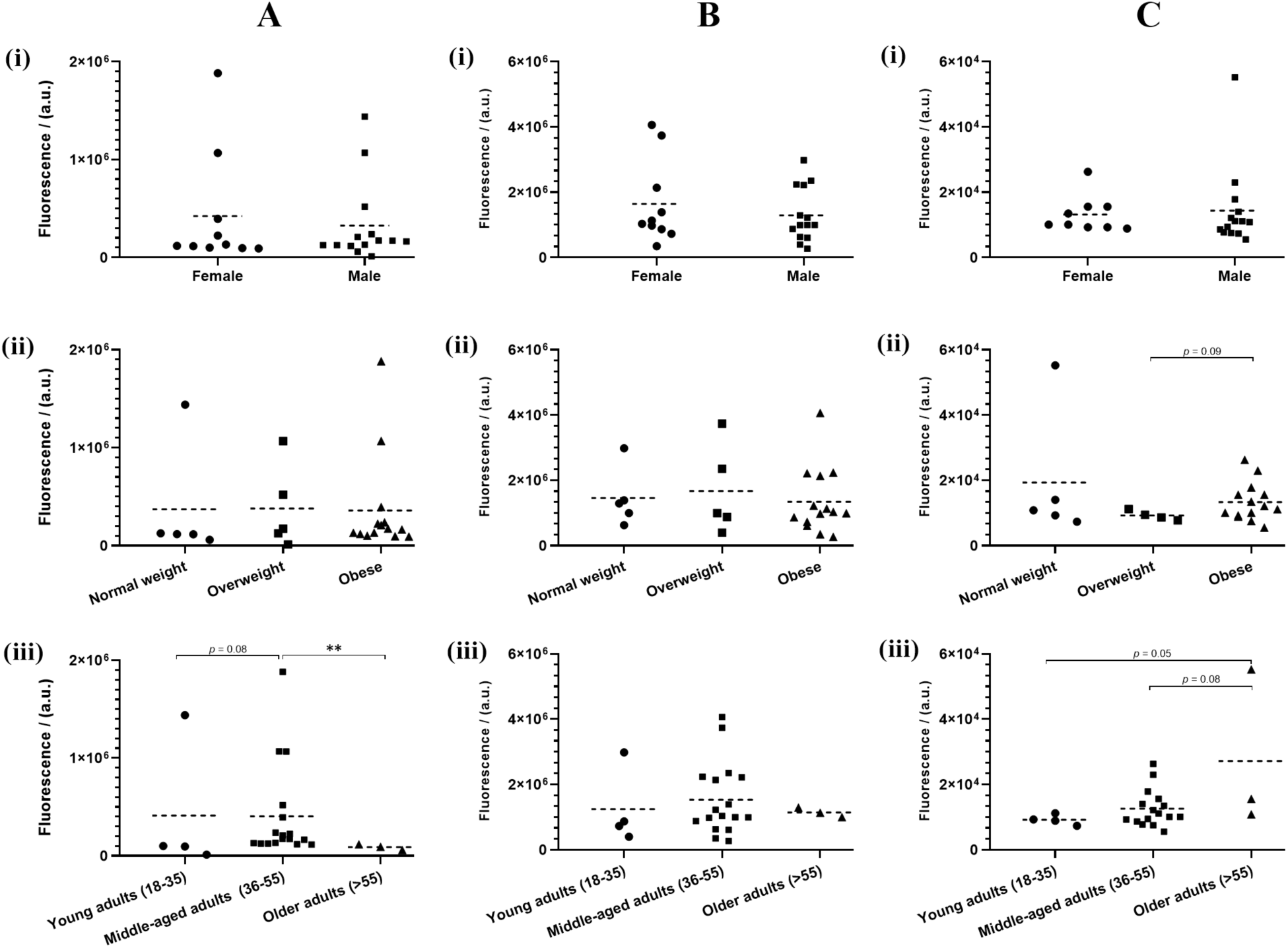
Cleavage of fluorogenic substrates Bz-FVR-AMC (thrombin-like enzymes, panel **A**), Boc-QAR-AMC (trypsin-like enzymes, panel **B**) and MCA-KPLGL-Dpa(DNP)-AR-NH2 (MMPs, panel **C**) by enzymes in patients’ 10% synovial fluids were compared against patients’ demographics, **(i)** sex, **(ii)** BMI and **(iii)** age. Mann-Whitney U test was utilized to compare any statistical significance between different groups, \**p* < 0.05 and \*\**p* < 0.01. Each data point on the scatter dot plot represents the mean ± SEM (*N* ≥ 3).

**Figure S3.**
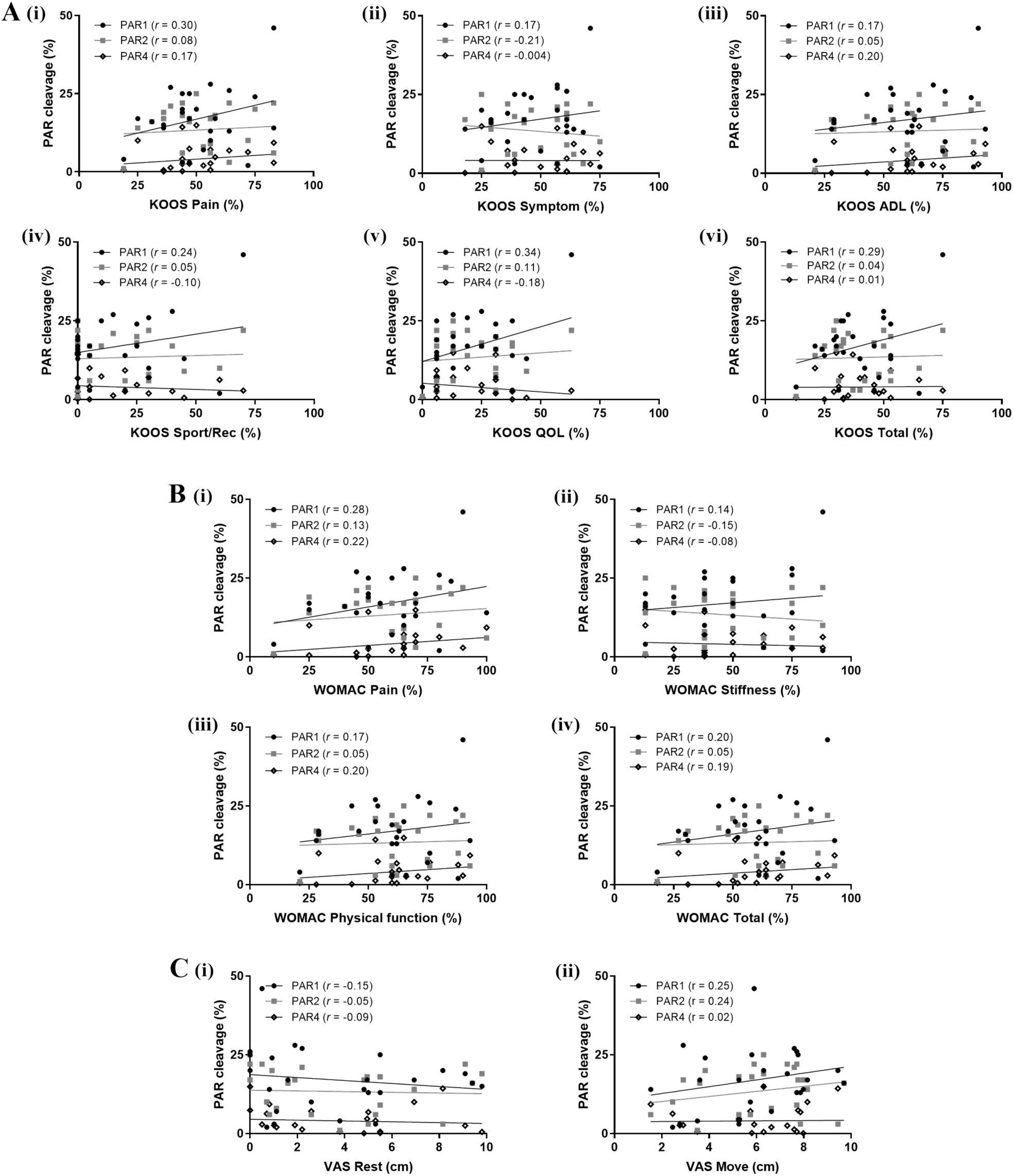
Correlation of the KOOS, WOMAC and VAS scores of twenty-five OA patients against the PAR1/2/4 activity. (**A**) KOOS, (**B**) WOMAC and (**C**) VAS against PARs. Pearson correlation coefficient analysis was performed to determine any correlation between PAR activity and the scores. Correlation coefficient interpretation was based on the following scale, *r* = ±1.00 to ±0.90: a very strong correlation, *r* = ±0.89 to ±0.70: a strong correlation, *r* = ±0.69 to ±0.40: a moderate correlation, *r* = ±0.39 to ±0.10: a weak correlation, and *r* = ±0.10 to ±0.00: negligible correlation.

**Table S1.** Statistical analysis of differences in cleavage of PAR1 by enzymes in human OA knee synovial fluids. Comparisons are made between healthy (Grade 0) and OA patients (Grade 1, 2 and 3) synovial fluids (10%) in nLuc-hPAR1-eYFP-CHO cells.

| **Table Analyzed**<br>**nLuc cleavage PAR1 vs Grade** |  |  |  |
| --- | --- | --- | --- |
| Column B | Grade 1 | Grade 2 | Grade 3 |
| vs. | vs. | vs. | vs. |
| Column A | Grade 0 | Grade 0 | Grade 0 |
| Mann Whitney test |  |  |  |
| P value | 0.1071 | 0.0741 | 0.3000 |
| Exact or approximate P value? | Exact | Exact | Exact |
| P value summary | ns | ns | ns |
| Significantly different ($P < 0.05$)? | No | No | No |
| Sum of ranks in column A,B | 18 , 60 | 18 , 73 | 21 , 45 |
| Mann-Whitney U | 8 | 8 | 11 |
| Difference between medians |  |  |  |
| Median of column A | 8.667, n=4 | 8.667, n=4 | 8.667, n=4 |
| Median of column B | 17.53, n=8 | 15.94, n=9 | 17.33, n=7 |
| Difference: Actual | 8.867 | 7.273 | 8.667 |
| Difference: Hodges-Lehmann | 7.817 | 7.650 | 7.055 |
| 96.64% CI of difference | -3.925 to 22.52 | -5.633 to 19.97 | -9.300 to 21.83 |
| Exact or approximate CI? | Exact | Exact | Exact |

**Table S2.** Statistical analysis of differences in cleavage of PAR2 by enzymes in human OA knee synovial fluids. Comparisons are made between healthy (Grade 0) and OA patients (Grade 1, 2 and 3) synovial fluids (10%) in nLuc-hPAR2-eYFP-CHO cells.

| **Table Analyzed**<br>**nLuc cleavage PAR2 vs Grade** |  |  |  |
| --- | --- | --- | --- |
| Column B | Grade 1 | Grade 2 | Grade 3 |
| vs. | vs. | vs. | vs. |
| Column A | Grade 0 | Grade 0 | Grade 0 |
| Mann Whitney test |  |  |  |
| P value | 0.1838 | 0.1650 | 0.3939 |
| Exact or approximate P value? | Exact | Exact | Exact |
| P value summary | ns | ns | ns |
| Significantly different ($P < 0.05$)? | No | No | No |
| Sum of ranks in column A,B | 32 , 46 | 35 , 56 | 26 , 40 |
| Mann-Whitney U | 10 | 11 | 12 |
| Difference between medians |  |  |  |
| Median of column A | 16.40, n=4 | 16.40, n=4 | 16.40, n=4 |
| Median of column B | 14.48, n=8 | 15.64, n=9 | 16.97, n=7 |
| Difference: Actual | -1.917 | -0.7600 | 0.5667 |
| Difference: Hodges-Lehmann | -4.683 | -5.867 | -4.783 |
| 96.64% CI of difference | -17.07 to 7.867 | -19.00 to 5.433 | -20.60 to 8.200 |
| Exact or approximate CI? | Exact | Exact | Exact |

**Table S3.**
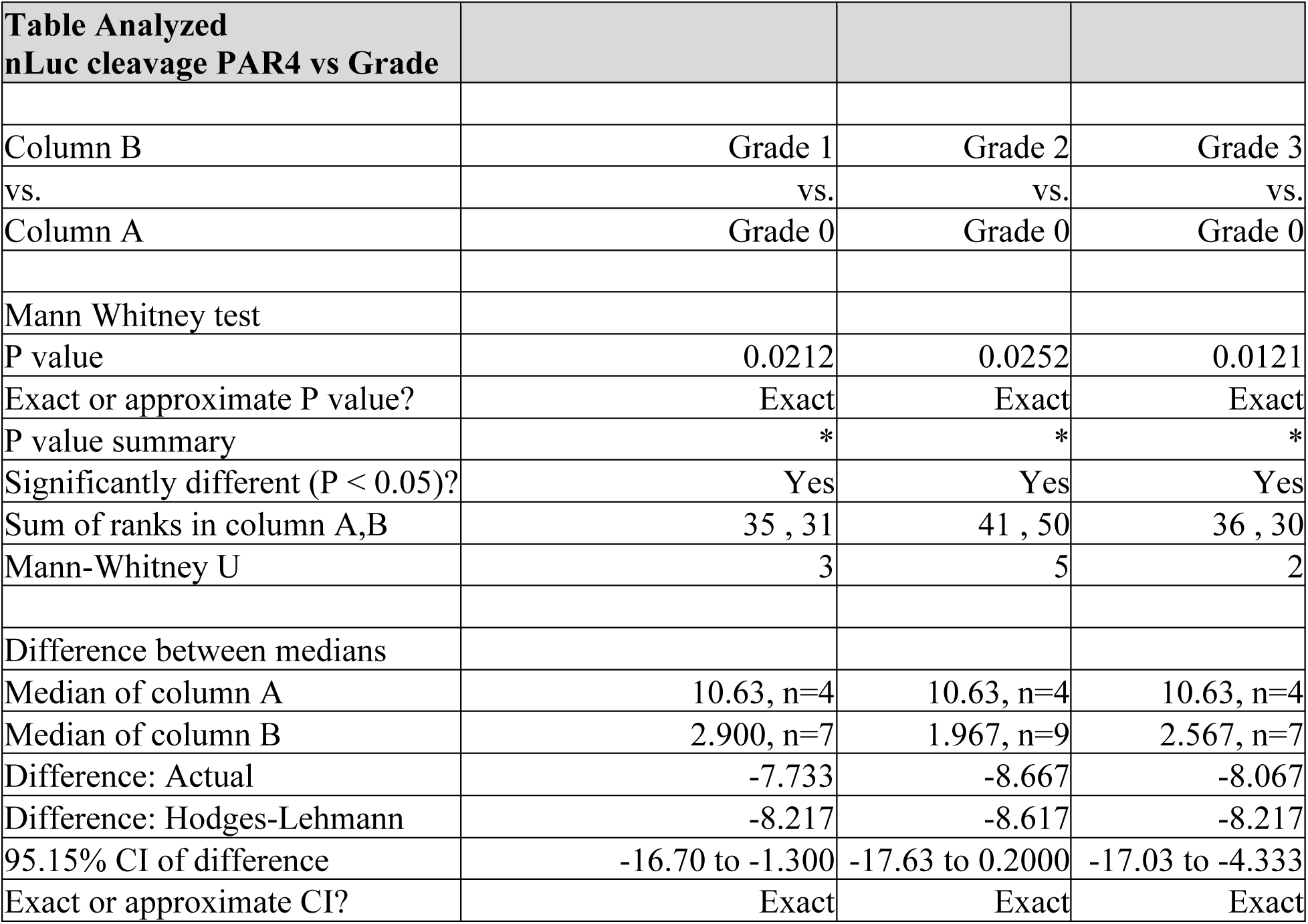
Statistical analysis of differences in cleavage of PAR4 by enzymes in human OA knee synovial fluids. Comparisons are made between healthy (Grade 0) and OA patients (Grade 1, 2 and 3) synovial fluids (10%) in nLuc-hPAR4-eYFP-CHO cells.

**Table S4.** KOOS, WOMAC and VAS scores of the OA patients.

| Patient | KOOS / (%) |  |  |  |  |  | WOMAC / (%) |  |  |  | VAS / (cm) |  |
| --- | --- | --- | --- | --- | --- | --- | --- | --- | --- | --- | --- | --- |
|  | Pain | Symptom | ADL | Sport/Rec | QOL | Total | Pain | Stiffness | Physical function | Total | Rest | Move |
| **1** | 83 | 71 | 90 | 70 | 63 | 75 | 90 | 88 | 90 | 90 | 0.5 | 5.9 |
| **2** | 44 | 57 | 53 | 0 | 31 | 37 | 53 | 50 | 38 | 51 | 8.2 | 9.4 |
| **3** | 56 | 61 | 60 | 45 | 44 | 53 | 65 | 50 | 60 | 60 | 5.5 | 7.7 |
| **4** | 64 | 61 | 76 | 30 | 19 | 50 | 80 | 75 | 76 | 77 | 0.0 | 7.7 |
| **5** | 36 | 61 | 62 | 0 | 6 | 33 | 25 | 38 | 62 | 52 | 9.8 | 6.3 |
| **6** | 72 | 75 | 88 | 60 | 31 | 65 | 88 | 80 | 88 | 86 | 0.7 | 2.5 |
| **7** | 56 | 36 | 76 | 30 | 13 | 42 | 65 | 38 | 76 | 71 | 2.6 | 5.7 |
| **8** | 75 | 46 | 87 | 25 | 31 | 53 | 85 | 50 | 87 | 83 | 0.9 | 3.8 |
| **9** | 83 | 64 | 93 | 20 | 6 | 53 | 93 | 100 | 75 | 93 | 0.8 | 1.5 |
| **10** | 31 | 29 | 29 | 0 | 31 | 24 | 40 | 13 | 29 | 30 | 9.4 | 9.7 |
| **11** | 47 | 43 | 54 | 10 | 6 | 32 | 54 | 60 | 50 | 55 | 0.0 | 7.8 |
| **12** | 44 | 39 | 43 | 0 | 38 | 33 | 50 | 38 | 43 | 44 | 5.5 | 5.8 |
| **13** | 50 | 25 | 65 | 0 | 13 | 30 | 70 | 13 | 65 | 61 | 0.0 | 6.3 |
| **14** | 64 | 68 | 62 | 0 | 6 | 40 | 62 | 70 | 63 | 64 | 5.0 | 7.9 |
| **15** | 53 | 50 | 75 | 30 | 31 | 48 | 60 | 38 | 75 | 69 | 1.1 | 6.6 |
| **16** | 19 | 25 | 21 | 0 | 0 | 13 | 10 | 13 | 21 | 18 | 3.8 | 3.5 |
| **17** | 47 | 43 | 46 | 0 | 13 | 30 | 55 | 50 | 46 | 48 | 1.6 | 3.6 |
| **18** | 56 | 57 | 71 | 40 | 25 | 50 | 65 | 75 | 71 | 70 | 1.9 | 2.9 |
| **19** | 44 | 36 | 60 | 0 | 6 | 29 | 50 | 25 | 60 | 55 | 9.1 | 7.3 |
| **20** | 39 | 57 | 53 | 15 | 13 | 35 | 45 | 38 | 53 | 50 | 2.2 | 7.6 |
| **21** | 58 | 61 | 63 | 25 | 25 | 46 | 63 | 70 | 50 | 64 | 5.0 | 5.3 |
| **22** | 47 | 57 | 66 | 20 | 38 | 46 | 50 | 75 | 66 | 64 | 1.0 | 2.8 |
| **23** | 36 | 18 | 28 | 5 | 38 | 25 | 45 | 25 | 28 | 31 | 4.8 | 8.0 |
| **24** | 44 | 39 | 60 | 5 | 6 | 31 | 65 | 63 | 60 | 61 | 5.3 | 5.3 |
| **25** | 25 | 29 | 29 | 5 | 19 | 21 | 29 | 25 | 13 | 27 | 6.9 | 8.2 |

KOOS, Knee Injury and Osteoarthritis Outcome Score; WOMAC, Western Ontario and McMaster Universities Osteoarthritis Index; VAS, Visual Analog Scale; ADL, Activities in Daily Living function; Sport/Rec, Sport and Recreation function; QOL, Quality of Life

